# The NKCC1 inhibitor bumetanide has no discernible effect on plasma cell survival, persistence or antibody secretion

**DOI:** 10.64898/2026.05.22.727109

**Authors:** Freeda Dsouza, David Tarlinton, Zhoujie Ding, Marcus James Robinson

**Affiliations:** Department of Immunology, Monash University, Melbourne, VIC, Australia

## Abstract

Long-lived plasma cells (LLPC) sustain humoral immunity but also contribute to the persistence of pathogenic autoantibodies in autoimmune diseases. New therapies targeting LLPC are therefore desirable. Recent studies have shown increased expression of *Slc12a2*, encoding the Na^+^ -K^+^ -Cl^−^ cotransporter (NKCC1), in LLPC. This study investigated whether NKCC1 activity was required for plasma cell survival, persistence or secretion of antibodies. Across *in vitro* and *in vivo* settings, mouse plasma cell survival was undiminished by treatment with the NKCC1 inhibitor bumetanide. Acute *in vivo* bumetanide treatment did not diminish plasma cell numbers, nor show any demonstrable impact on the survival of phenotypically mature I-A/I-E^lo^SLAMF6^lo^ plasma cells. With genetic plasma cell timestamping, even the survival of persistent LLPC was unaffected by bumetanide. Plasma cell secretory capacity, assessed by measuring IgM and IgG2b secretion in culture over three days, was also unaltered by bumetanide. Overall, these results show that pharmacological inhibition of NKCC1 is not sufficient to impair plasma cell survival, persistence or antibody secretion. Despite elevated *Slc12a2* mRNA expression in LLPC, NKCC1 alone does not represent a critical plasma cell survival pathway, highlighting the resilience of plasma cells and the challenges associated with therapeutically targeting LLPC.

## Introduction

Bumetanide is a loop diuretic used in the treatment of oedema associated with congestive heart failure, renal and hepatic conditions, acute pulmonary congestion and premenstrual syndrome. It produces dose-related diuresis with urinary excretion of sodium and chloride over a dose range of 0.5-3 mg in humans [1]. Loop diuretics primarily act in the thick ascending limb of the loop of Henle in the kidney by inhibiting Na^+^, K^+^ and Cl^−^ reabsorption, consequently reducing the medullary concentration gradient and water reabsorption and increasing salt and water excretion, along with potassium loss [2]. Bumetanide specifically inhibits this transporter in the ascending limb and can also act on the proximal tubules [1]. Elimination occurs via renal, hepatic, or biliary routes with a half-life of 1-1.5 hours [1].

Beyond its established diuretic use, bumetanide has been widely used experimentally as an inhibitor of NKCC1. In mice, at 25 mg/kg it has been shown to preserve blood-brain barrier (BBB) integrity and improve functional outcomes following traumatic brain injury, supporting the biological activity of bumetanide-mediated NKCC1 inhibition [3]. Bumetanide has also been investigated in neurological disorders, where NKCC1 contributes to intracellular chloride accumulation and altered neuronal excitability [4]. In neurons, the balance between the chloride importer NKCC1 and the exporter KCC2 determines the inhibitory effect of GABA [5]. Reduced NKCC1 activity driven by bumetanide treatment can lower intracellular chloride and enhance GABA-mediated inhibitory signaling, making NKCC1 inhibition a possible therapeutic strategy in conditions such as epilepsy, neonatal asphyxia, traumatic brain injury, autism, and chronic pain [6].

Recent studies have found expression of the gene encoding the NKCC1 cotransporter in plasma cells (PC), suggesting a potential role for ion transport in maintaining homeostasis and survival [7]. Notably, among all PC, the oldest cells were the highest expressing, raising the possibility that targeting NKCC1 using bumetanide could influence PC persistence. However, direct evidence demonstrating that NKCC1 inhibition compromises PC survival or function remains absent to the best of our knowledge. Here, we sought to establish if LLPC rely on NKCC1 for survival, persistence or Immunoglobulin secretion using Bumetanide to inhibit the receptor on PC *in vitro* and *in vivo*.

## METHODS

### ANIMALS

C57BL/6, BLTcre [8] and BLT_Cre_ .Mcl^fl/+^ [9] mice were bred and housed in the specific pathogen-free facilities at the Monash University Animal Research Platform (MARP), Melbourne, Australia. Mice were group housed in a controlled environment maintained on a 12-hour light/dark cycle, with *ad libitum* access to water and a standard rodent maintenance diet (Barastoc Rat and Mouse Diet, Ridley AgriProducts, Australia). Cages were equipped with nesting material, such as shredded paper or tissues, and environmental enrichment was provided on a rotating basis, including egg cartons, tunnels, aspen sticks and Eppendorf tubes. All experiments were conducted with the approval of the Alfred Research Alliance Animal Ethics Committee. Both male and female mice were used in these experiments.

### PREPARATION OF MOUSE BONE MARROW SINGLE-CELL SUSPENSION

To obtain PC from murine bone marrow, mice were euthanised by CO_2_ asphyxiation and immediately subjected to cervical dislocation as a secondary method before tissue collection. Femurs and spines were dissected and surrounding muscle and connective tissues removed. Bones were mechanically disrupted using a mortar and pestle in a buffer containing 0.5% bovine serum albumin (BSA, pH 7.2), 2 mM ethylenediaminetetraacetic acid (EDTA, pH 8) in Dulbecco’s phosphate-buffered saline (PBS, pH 7). The resulting cell suspension was pressed through a 70 µm cell strainer using the rubber end of a 3 mL syringe. Red blood cells were lysed using red cell lysis buffer (156 mM NH_4_Cl, 11.9 mM NaHCO_3_, 97 µM EDTA, pH 7.3), after which remaining cells were resuspended in 0.5% BSA/PBS and centrifuged at 350 *g* for 4 minutes. Cells were then strained through 50 µm nylon gauze. For *in vitro* cultures, cells were then counted and seeded at 5 × 10^6^ cells per well in 500 µL or RPMI-1640 supplemented with 10% FCS (pH 7.2), 55 µM 2-mercaptoethanol (pH 7.1), 10 mM HEPES (pH 7.4), 1 mM sodium pyruvate (pH 7), and 1% penicillin-streptomycin solution (pH 6.5). Seeded cells were kept on ice before staining. Cell counts were obtained using a CellDrop Automated Cell Counter (DeNovix).

### FLOW CYTOMETRY

Following cell counting for *ex vivo* analyses, cells were resuspended in flow buffer (0.5 % BSA, 2 mM EDTA in PBS) and incubated with a surface stain containing 1.25 µg/mL 2.4G2 antibody (WEHI Monoclonal Antibody Facility) and normal rat serum (1% v/v) for 5 minutes. This was followed by staining with antibodies against the surface markers SLAMF6, I-A/I-E, CD98, IgD and efluor780 as a live/dead stain, and the cells were incubated on ice for 30 minutes. After incubation, cells were washed twice with flow buffer and centrifuged at 350 g for 4 minutes. The washed cells were then resuspended in flow buffer and analysed on a BD LSR Fortessa X20 and analysed in FlowJo Software (Treestar).

### BUMETANIDE PREPARATION FOR IN VITRO USE

Bumetanide (Sigma-Aldrich, St Louis, MO, USA) was dissolved in 100 % dimethyl sulfoxide (DMSO, pH 7) to prepare a 20 mM stock solution, which was then diluted to 20 µM using cRPMI (pH 7) and serially diluted in vehicle for culture tests.

### INTRAPERITONEAL BUMETANIDE ADMINISTRATION

Bumetanide was dissolved in PBS at 0.75 mg/mL and administered to mice via intraperitoneal injection at 25 mg/kg once every six hours three times on a single day. Mice receiving PBS alone were used as controls.

### TAMOXIFEN ADMINISTRATION

Tamoxifen (Sigma-Aldrich, St Louis, MO, USA) was administered by oral gavage at a dose of 200 mg/kg, using a 60 mg/mL formulation in 90% (v/v) peanut oil and 10% (v/v) ethanol.

### ELISA (ENZYME-LINKED IMMUNOSORBENT ASSAY)

Isolated bone marrow cells were seeded at 5 × 10^6^ cells per well and incubated for 72 hours at 37°C humidified in air with 5% CO_2_. Culture supernatants were then collected and stored at -20 °C until analysis. For ELISA, 96-well EIA/RIA plates (Corning, Glendale, AZ, USA) were coated overnight at 4°C with 2 µg/mL goat-anti-mouse IgG2b or goat-anti-mouse IgM (Southern Biotech) diluted in PBS. Plates were washed, and culture supernatants were added in 5-fold serial dilutions from neat in blocking buffer containing 1% fetal bovine serum (FBS), 0.6% (w/v) skim milk powder and 0.05% v/v Tween-20. Plates were incubated for 2 hours at room temperature and then washed before addition of HRP-conjugated detection antibodies (anti-IgG2b-HRP and anti-IgM-HRP; Southern Biotech) for 1 hour at room temperature. After washing, TMB substrate (Thermo Fisher Scientific) was added and incubated in the dark for colour development. The reaction was stopped using 2 M HCl, and absorbance was measured at 450 nm using a microplate reader (Thermo Fisher Scientific). Immunoglobulin concentrations were determined by comparison to standard curves generated using simple linear regression restricted to the linear dynamic range of the assay.

### STATISTICS

Flow data were analysed and exemplars exported from FlowJo Software, while other data were graphed and analysed statistically using RStudio (Posit). *P* < 0.05 was considered statistically significant throughout.

## Results

### Bumetanide does not alter plasma cell survival *in vitro*

We examined the *in vitro* effects of bumetanide on PC survival, with a particular focus on I-A/I-E^lo^SLAMF6^lo^ PC, a state associated with PC longevity and with high *Slc12a2* (NKCC1) transcription [7], to assess whether NKCC1 activity contributes to the survival of PC. Five million total bone marrow cells were cultured with Bumetanide at various concentrations or in media containing DMSO only as control, for 6 or 24 hours. PC were identified by flow cytometry, and enumerated by determining their representation from the total live singlet gate and multiplying that frequency by the total cell count, as determined by Celldrop.

At 6 hours, PC numbers were equivalent across all concentrations (Figure 1A), and bumetanide did not alter the subset distribution (*P* > 0.05 for all comparisons, Figure 1B). At 24 hours, PC numbers were similar across Bumetanide concentrations, and again, no change in subset representation across doses (*P* > 0.05 for all comparisons, Figure 1D). Importantly, the frequency of I-A/I-E^lo^SLAMF6^lo^ PC, those predicted to express the highest *Slc12a2* amounts, was unchanged across all treatment conditions, as were other PC subsets (Figure 1D). Overall, these findings indicate that bumetanide does not impair PC survival *in vitro* and does not affect the representation of I-A/I-E^lo^SLAMF6^lo^ PC.

**Figure 1:**
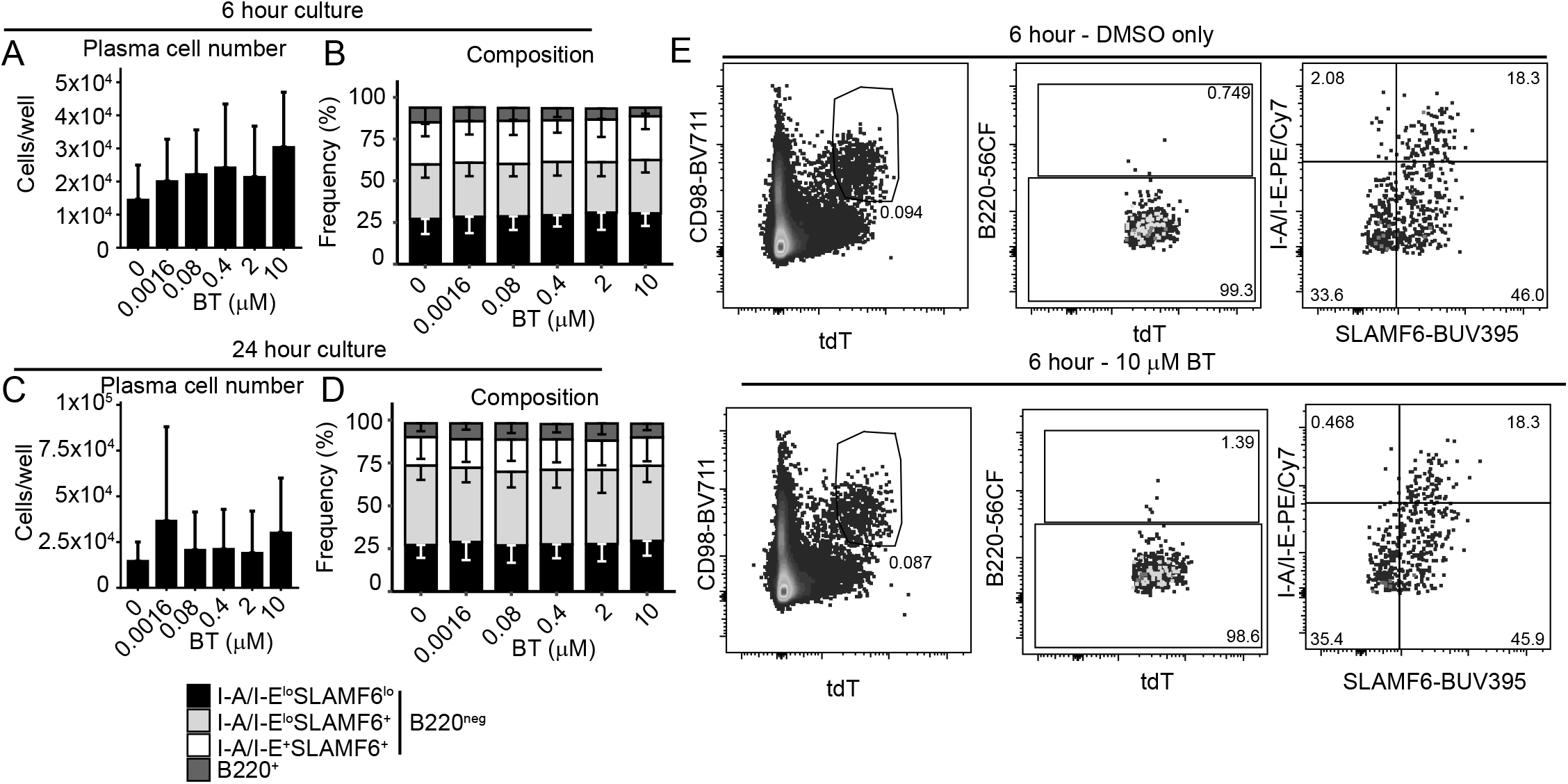
Bumetanide does not impair PC survival *in vitro*. Bone marrow cells were cultured with bumetanide (10 µM, 2 µM, 0.4 µM, 0.08 µM, 0.016 µM) or DMSO control for 6 and 24 hours. PC were analysed by flow cytometry and identified as CD98^+^tdT^+^ live singlet cells. At six hours (A) total PC counts and (B) subset representation was determined, similarly at 24 hours (C) total PC counts and (D) subset representation was determined. Statistical significance was assessed using (A, C) repeated measures one-way ANOVA with Sidak’s multiple comparisons test or (B, D) repeated-measures two-way ANOVA with Sidak’s multiple comparisons test. Data represent arithmetic mean ± SD of n=8 (A, B) or n=4 (C, D) independent biological replicates combined from eight (A, B) or four (C, D) separate experiments.

### Acute bumetanide treatment does not alter plasma cell subset representation or number *in vivo*

To determine whether NKCC1 supports PC survival *in vivo*, particularly I-A/I-E^lo^SLAMF6^lo^ PC, mice were treated with bumetanide. Mice received intraperitoneal injections of bumetanide in PBS or PBS alone three times in a single day, 6 hours apart. Bone marrow cells were collected 24 hours after the first dose and analyzed by flow cytometry and cell counting.

PC and the subsets were readily identifiable in both PBS- and Bumetanide-treated mice (Figure 2A). Total PC numbers were comparable between PBS and bumetanide-treated mice (Figure 2B). Analysis of PC subset frequencies revealed a treatment-associated change in subset representation, but surprisingly this was revealed as an increased representation of I-A/I-E^lo^SLAMF6^lo^ PC, rather than the specific loss of these cells, following bumetanide treatment (Figure 2C). Importantly, the absolute numbers of I-A/I-E^lo^SLAMF6^lo^ PC, representing the subset with the highest *Slc12a2* expression, were unchanged following bumetanide treatment (Figure 2D).

**Figure 2:**
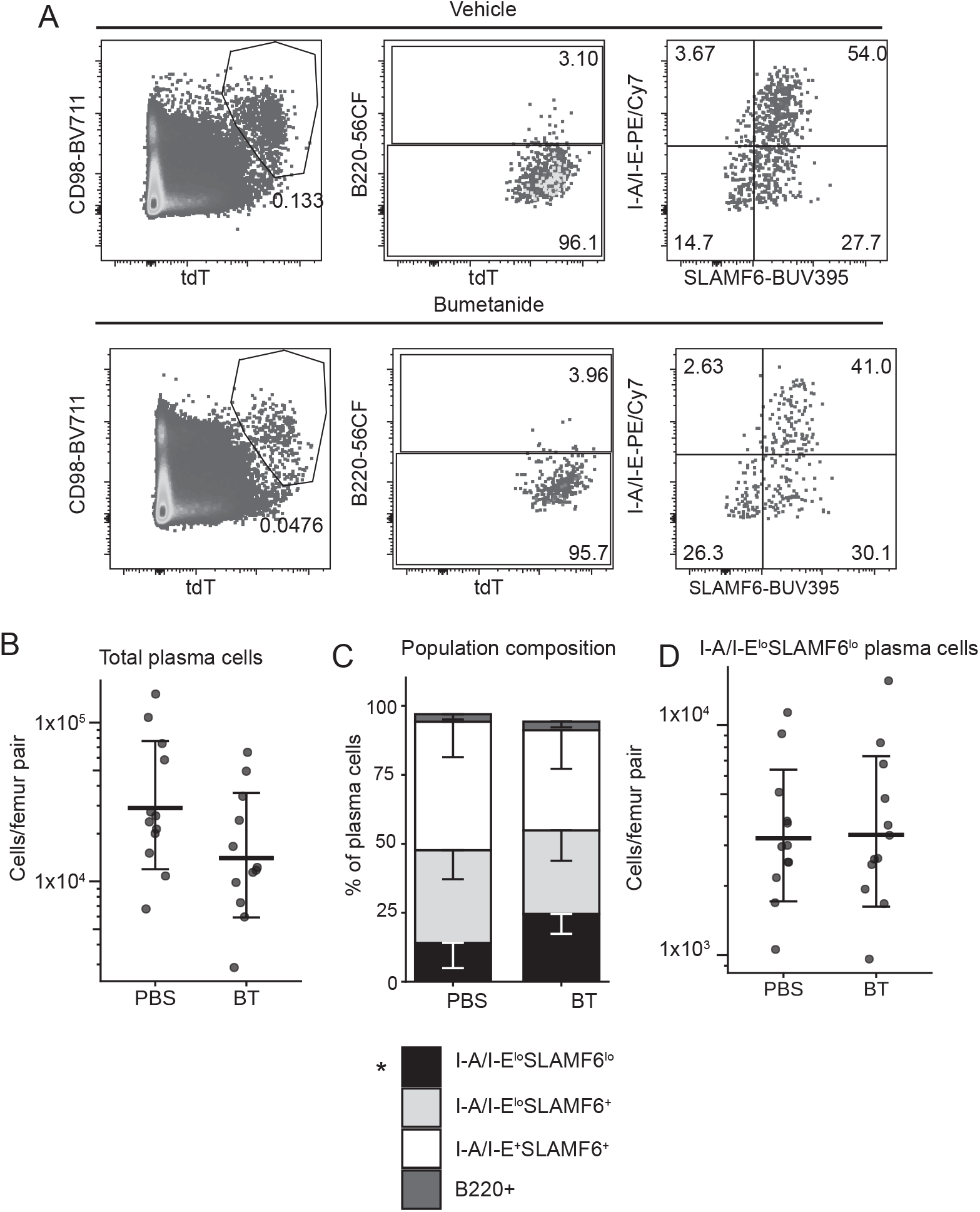
Acute *in vivo* bumetanide treatment does not alter PC subsets or absolute PC numbers. Mice were injected intraperitoneally with bumetanide in PBS, or with PBS alone three times over 24 hours, with injections administered six hours apart. Femurs were harvested 24 hours after the first injection, and total bone marrow cells were analysed by flow cytometry. (A) PC were identified as CD98^+^tdT^+^ events among live IgD^lo^ singlet cells, and compartmentalized into B220^+^ and B220^neg^ subsets, then further subsetted by I-A/I-E and SLAMF6. (B) Absolute PC numbers (C) subset representation, and (D) number of I-A/I-E^lo^SLAMF6^lo^ PC. Statistical significance was assessed using (B, D) Unpaired Student’s t-test after log-transformation or (C) two-way repeated-measures ANOVA with Sidak’s multiple comparisons test. (B, D) Data symbols show individual mice and lines show geometric mean ± geometric SD factor or (C) arithmetic mean – SD of n=12 independent biological replicates per group, combined from two experiments with n=6 mice per group per experiment. * P < 0.05, PBS vs BT. BT = bumetanide.

These findings therefore indicate that acute NKCC1 inhibition does not impair PC survival *in vivo* over 24 hours and does not reduce the representation of I-A/I-E^lo^SLAMF6^lo^ PC.

### NKCC1 inhibition does not reduce established LLPC *in vivo*

PC survival and persistence represent different features of the PC program [10] and we therefore wanted to examine whether NKCC1 function affected PC persistence. PC were therefore time-stamped using a tamoxifen-inducible human CD4 reporter system. Mice were administered tamoxifen on day 0 to genetically label PC, then received intraperitoneal injections of bumetanide or PBS on day 18, three times with injections administered six hours apart for a transient treatment window, followed by resting the mice for ten further days, allowing for any chronic impact of the transient treatment to play out, and bone marrow was harvested on day 28, ten days after bumetanide exposure (Figure 3A). The cells were analysed by flow cytometry to assess total hCD4^+^ PC, particularly the I-A/I-E^lo^SLAMF6^lo^ PC, and overall subset distribution.

**Figure 3:**
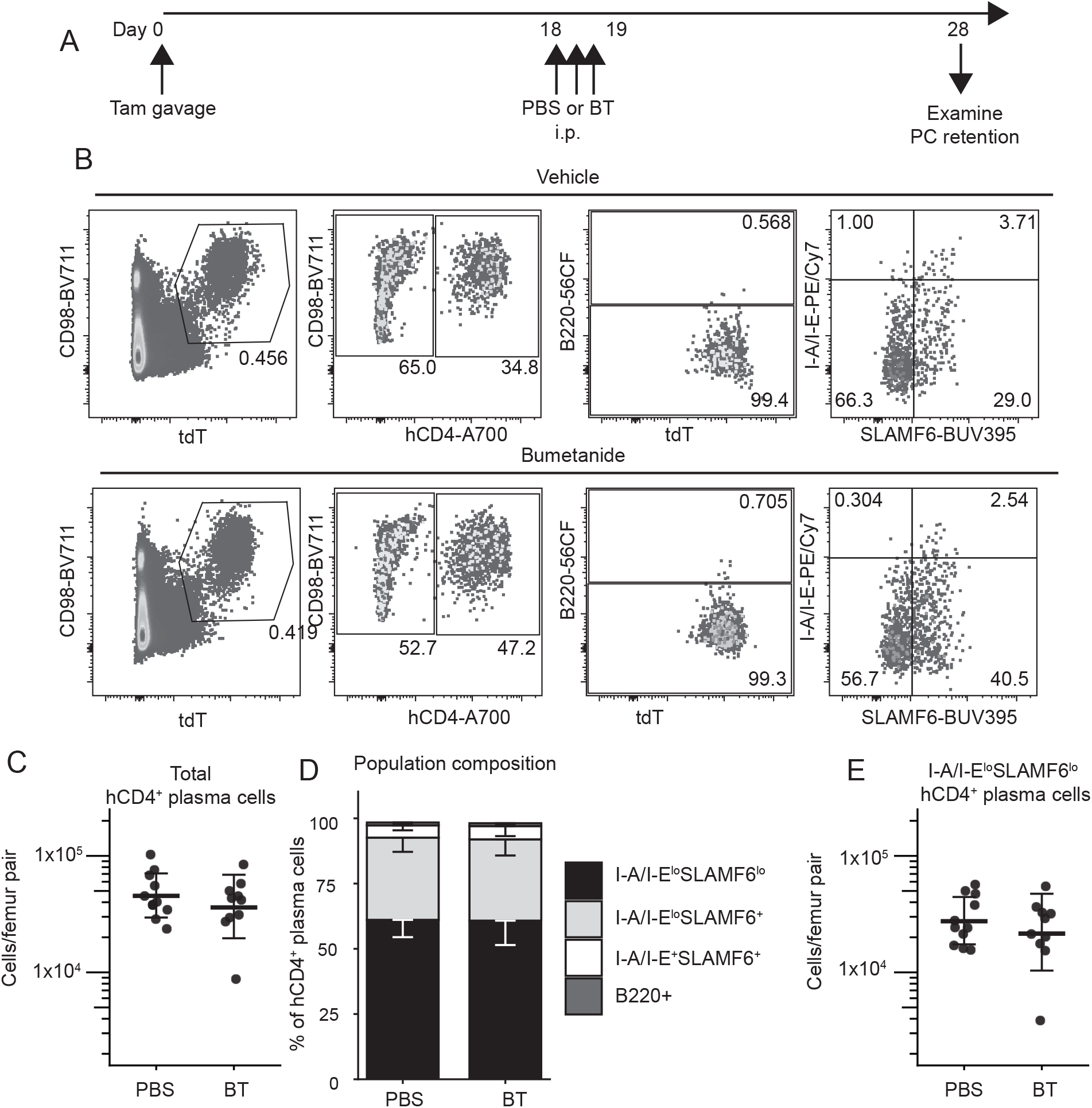
Transient NKCC1 inhibition does not alter PC persistence. (A) Schematic showing that mice were gavaged with Tamoxifen to timestamp PC, and 18 days later, i.p. administered Bumetanide or PBS three times, six hours apart and returned to stock housing. Ten days later bone marrow was harvested and (B) by flow cytometry, assessed for persistent PC as CD98^+^tdT^+^hCD4^+^ cells among live IgD^lo^ singlet cells, and compartmentalized into B220^+^ and B220^neg^ subsets, then further subsetted by I-A/I-E and SLAMF6. (C) Absolute hCD4^+^ PC numbers, (D) subset representation, and (E) number of hCD4^+^ I-A/I-E^lo^SLAMF6^lo^ PC in the femurs. Statistical significance was assessed using (C, E) Unpaired Student’s t-test after log-transformation or (D) two-way repeated-measures ANOVA with Sidak’s multiple comparisons test. Data symbols show (C, E) individual mice and lines show geometric mean ± geometric SD factor, or (D) mean + SD of n=10-11 mice per treatment combined from two separate experiments. BT = bumetanide.

hCD4^+^ PC were identified and split into subsets (Figure 3B) and found to be similarly abundant in PBS and bumetanide-treated mice (Figure 3C). The population composition was similar across treatments (Figure 3D), such that the absolute number of I-A/I-E^lo^SLAMF6^lo^ hCD4+ PC, representing the matured PC subset, were unchanged following bumetanide treatment (Figure 3E).

Collectively, these findings show that NKCC1 inhibition does not reduce the survival of established LLPC *in vivo*, further supporting the conclusion that NKCC1 activity is not required for the maintenance of the LLPC.

### NKCC1 inhibition does not modulate IgM or IgG2b secretion

Although PC survival was not dependent on NKCC1 activity, it remained possible that NKCC1 inhibition could affect PC function. Therefore, to determine whether NKCC1 inhibition affected antibody secretion, total IgM and IgG2b secretion were measured in culture supernatants from bone marrow cells following treatment with either DMSO vehicle or 10 µM bumetanide for 72 hours. Measuring both an unswitched isotype, IgM, and a class-switched isotype, IgG2b, enabled assessment of whether NKCC1 inhibition broadly affects antibody secretion. Analysis of IgM secretion revealed no significant difference between treatment groups (paired Student’s t-test *P* = 0.2547; Figure 4A), nor did IgG2b secretion differ (*P* = 0.0804, Figure 4B). We conclude that pharmacological inhibition of NKCC1 with bumetanide does not impair IgM or IgG2b antibody secretion by bone marrow PC.

**Figure 4:**
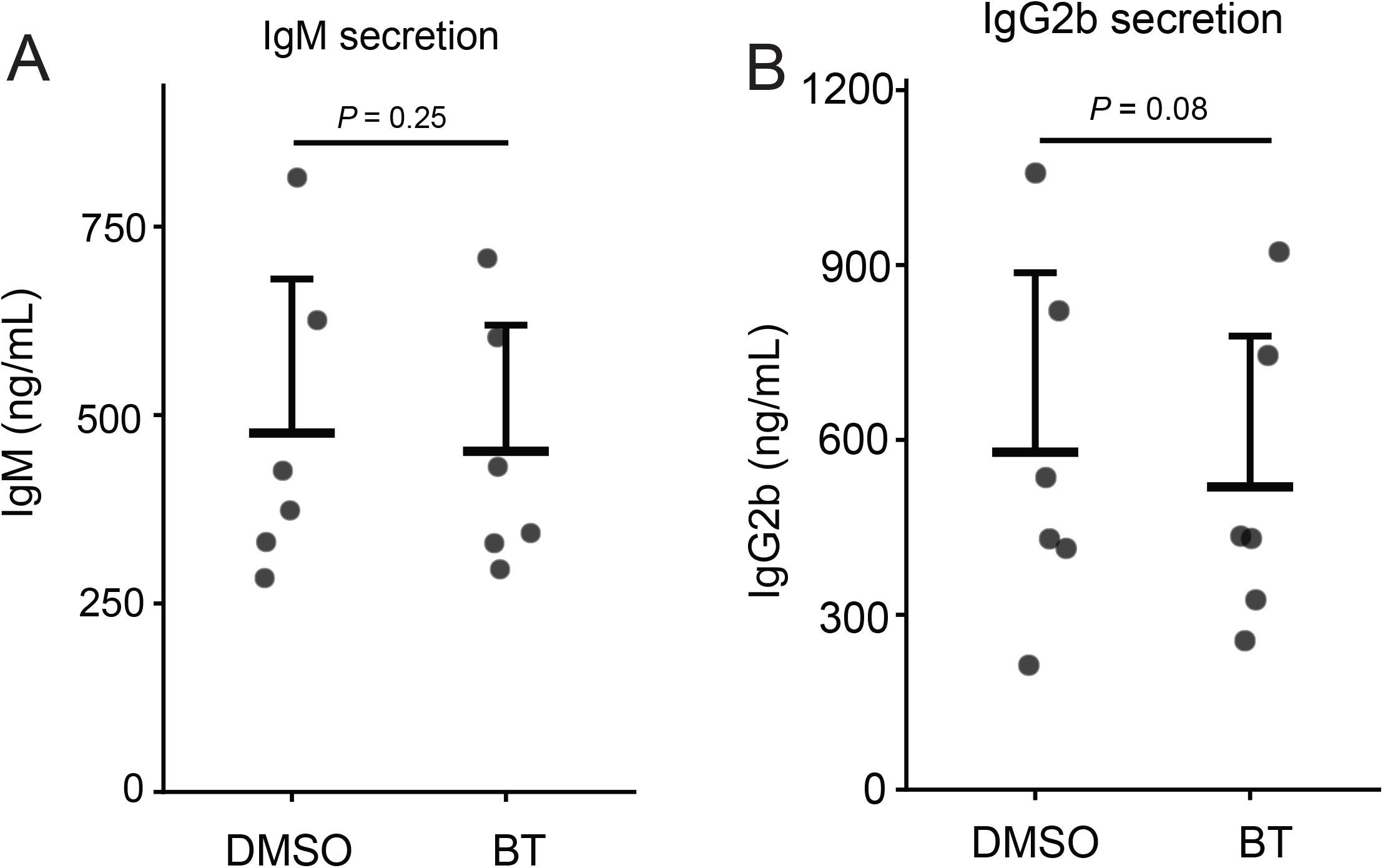
NKCC1 inhibition does not impair *in vitro* antibody secretion. Bone marrow cells were cultured for 72 hours in the presence of 10 μM bumetanide or 0.1% DMSO vehicle control. Culture supernatants were collected, and total (A) IgM and (B) IgG2b concentrations were quantified using ELISA. Statistical significance was assessed using a paired Student’s t-test. Data lines show arithmetic mean ± SD and symbols individual replicates of n=6 independent biological replicates combined from two separate experiments with n=3 individual biological replicates per experiment.

## Discussion

This study aimed to determine whether inhibition of the NKCC1 cotransporter affects PC survival or function, with particular focus on LLPC that exhibit elevated *Slc12a2* mRNA expression. Across *in vitro*, acute *in vivo* and long-term *in vivo* models, pharmacological inhibition using bumetanide did not significantly impair PC survival. Total PC numbers remained relatively stable following treatment, and the I-A/I-E^lo^SLAMF6^lo^ PC subset was preserved under all experimental conditions. Together, these findings indicate that NKCC1 inhibition alone is insufficient to impair PC survival.

*in vitro* exposure of PC to bumetanide across a range of concentrations did not alter total PC numbers at either 6 hours or 24 hours, indicating that acute NKCC1 inhibition does not directly induce PC death. The concentration range used in this study was selected based on previous *in vitro* studies demonstrating effective NKCC1 inhibition within similar micromolar ranges in a variety of cell types, including erythrocytes and a macrophage cell line [11-14]. Based on these findings, an effect on PC survival might have been expected if NKCC1 activity were essential for maintaining viability. However, PC numbers remained stable across all treatment conditions, showing that bumetanide alone does not substantially affect PC survival under the conditions tested. Further, importantly, the I-A/I-E^lo^SLAMF6^lo^ PC subset remained stable across all treatment conditions, indicating that cells enriched for aged PC, which have the highest *Slc12a2* expression, are not particularly sensitive to NKCC1 inhibition *in vitro*.

Consistent with the *in vitro* findings, acute *in vivo* administration of bumetanide over 24 hours did not reduce total PC numbers or the absolute number of I-A/I-E^lo^SLAMF6^lo^ PC within the bone marrow. I-A/I-E^lo^SLAMF6^lo^ PC represent a more mature, PC population enriched for LLPC which exhibit the highest *Slc12a2* expression. Although an increased representation of I-A/I-E^lo^SLAMF6^lo^ PC was observed following treatment, this occurred without any change in absolute PC numbers, suggesting proportional variation in subset distribution rather than a survival effect. These results demonstrate that short-term systemic NKCC1 inhibition does not deplete PC and does not selectively disadvantage the most mature PC.

The dose of bumetanide used in this study (25 mg/kg) was selected based on previous *in vivo* studies demonstrating biological activity at similar concentrations with a similar dosing regimen [3]; in mouse models of neurological injury, repeated administration of bumetanide at 25 mg/kg has been shown to produce protective effects associated with NKCC1 inhibition, indicating that this dose range is pharmacologically active *in vivo* [3]. Other rodent studies investigating NKCC1 function have also applied comparable doses of bumetanide and reported measurable physiological responses [15]. These findings support that the dosing strategy used in this study was likely sufficient to achieve systemic inhibition of NKCC1. Despite this, PC numbers were unchanged by treatment, indicating that NKCC1 inhibition at biologically active concentrations does not substantially affect PC survival *in vivo*.

To determine whether bumetanide exposure affects LLPC, a long-term lineage tracing *in vivo* model incorporating tamoxifen-induced human CD4 timestamping of the LLPC was used. This approach enabled unequivocal identification and tracking of the LLPC. Despite the select focus on LLPC, which have the highest transcription of *Slc12a2*, bumetanide did not reduce their numbers. PC subset frequencies remained comparable between the treatment groups, with statistical analysis failing to reveal any treatment-specific effects. These findings show that NKCC1 inhibition does not affect the persistence of LLPC.

In addition to assessing survival, PC function was evaluated by measuring IgM and IgG2b secretion following bumetanide treatment. IgM and IgG2b were selected as representative immunoglobulin classes that are produced by PC, allowing reliable quantification of total antibody secretion in culture. Antibody secretion, as quantified by ELISA, remained unchanged at the concentration of bumetanide tested. Given that PC are defined by their capacity to produce large quantities of antibodies, these findings indicate that NKCC1 inhibition does not impair the functional output of PC, at least under the conditions examined. Antibody secretion is a highly metabolically demanding process requiring a regulated ionic and osmotic balance. Therefore, if NKCC1 was critically required for maintaining intracellular homeostasis, a reduction in immunoglobulin production would have been expected.

Overall, the consistent preservation of PC numbers across both *in vitro* and *in vivo* models indicates that NKCC1 activity is not essential for PC survival. The absence of any significant change in IgM or IgG2b secretion further supports the conclusion that NKCC1 is not a primary regulator of PC viability or secretory capacity, at least in isolation. This does not rule out that the transporter is important, as it is possible that its function is compensated for by other ion transporters. Equally, although *Slc12a2* expression is elevated in LLPC [7], increased expression does not necessarily mean that PC depend on NKCC1 for their survival. Instead, NKCC1 may play a supportive role in PC physiology. The stability of PC despite NKCC1 inhibition suggests that alternative pathways are sufficient to maintain LLPC survival and function.

